# First chromosome-level genome assembly of a ribbon worm from the Hoplonemertea clade, *Emplectonema gracile*, and its structural annotation

**DOI:** 10.1101/2024.02.16.580704

**Authors:** Alberto Valero-Gracia, Nickellaus G. Roberts, Meghan Yap-Chiongco, Ana Teresa Capucho, Kevin M. Kocot, Michael Matschiner, Torsten H. Struck

**Affiliations:** Natural History Museum, University of Oslo, Blindern, P.O. Box 1172, 0318 Oslo, Norway; Department of Biological Sciences, University of Alabama, Tuscaloosa, AL 35487, USA; Alabama Museum of Natural History, University of Alabama, Tuscaloosa, AL 35487, USA

**Author notes:** Correspondence: Alberto Valero-Gracia, Natural History Museum, University of Oslo, Norway.

**Keywords:** de novo assembly, genome sequence, 3D genomics, Hi-C, HiFi

## Abstract

Genome-wide information has so far been unavailable for ribbon worms of the clade Hoplonemertea, the most species-rich class within the phylum Nemertea. While species within Pilidiophora, the sister clade of Hoplonemertea, possess a pilidium larval stage and lack stylets on their proboscis, Hoplonemertea species have a planuliform larva and are armed with stylets employed for the injection of toxins into their prey. To further compare these developmental, physiological, and behavioral differences from a genomic perspective, the availability of a reference genome of a Hoplonemertea species is crucial. To this end, we herein present the annotated chromosome-level genome assembly for *Emplectonema gracile* (Nemertea; Hoplonemertea; Monostilifera; Emplectonematidae), an easily collected nemertean well-suited for laboratory experimentation. The genome is 157.9 Mbp in span. Hi-C scaffolding yielded 15 putative chromosomes with a scaffold N50 of 10.0 Mbp and a BUSCO completeness score of 95.3%. Structural annotation predicted 20,684 protein-coding genes. The high-quality reference genome reaches an Earth BioGenome standard level of 7.C.Q50. These data will be highly useful for future investigations towards a better understanding of the evolution, development, morphology, and toxicology of Nemertea.

**Significance:** The genome of *Emplectonema gracile* is highly contiguous, well annotated, and shorter than those of the other two ribbon worm species sequenced to date. This genome is a valuable resource for studies on molecular ecology, venom evolution, and regeneration in marine invertebrates.

## Introduction

Nemerteans, commonly known as ribbon worms, are a phylum of about 1,200 species of predatory worms that exhibit spiral cleavage and a variety of life histories, typically including pelagic and benthic stages (Gibson 1994, Maslakova and Hiebert 2015). While nemerteans are mainly marine, some species have entered freshwater habitats, and a few have colonized moist, terrestrial habitats (Gibson 1994). Phylogenetically, Nemertea is nested within Spiralia (*sensu* Giribet and Edgecombe 2020). However, their exact phylogenetic position is not well established (Struck and Fisse 2008; Struck et al. 2014; Andrade et al. 2014; Laumer et al. 2015; Kocot et al. 2017; Bleidorn 2019; Marlètaz et al. 2019; Dràbková et al. 2022).

Nemertea is divided into three main clades: Paleonemertea, Pilidiophora, and Hoplonemertea (Figure 1A) (Andrade et al. 2014; Kvist et al. 2014). To date, only two nemertean nuclear genomes are available, both coming from species of the family Lineidae (Pilidiophora). One of these nemertean genomes, *Lineus longissimus*, meets the current Earth BioGenome Project standards for reference genomes (Kwiatkowski et al. 2021), while the second, the genome of *Notospermus geniculatus*, was published only at the level of scaffolds (Luo et al. 2018) (Figure 1A).

**Fig. 1.**
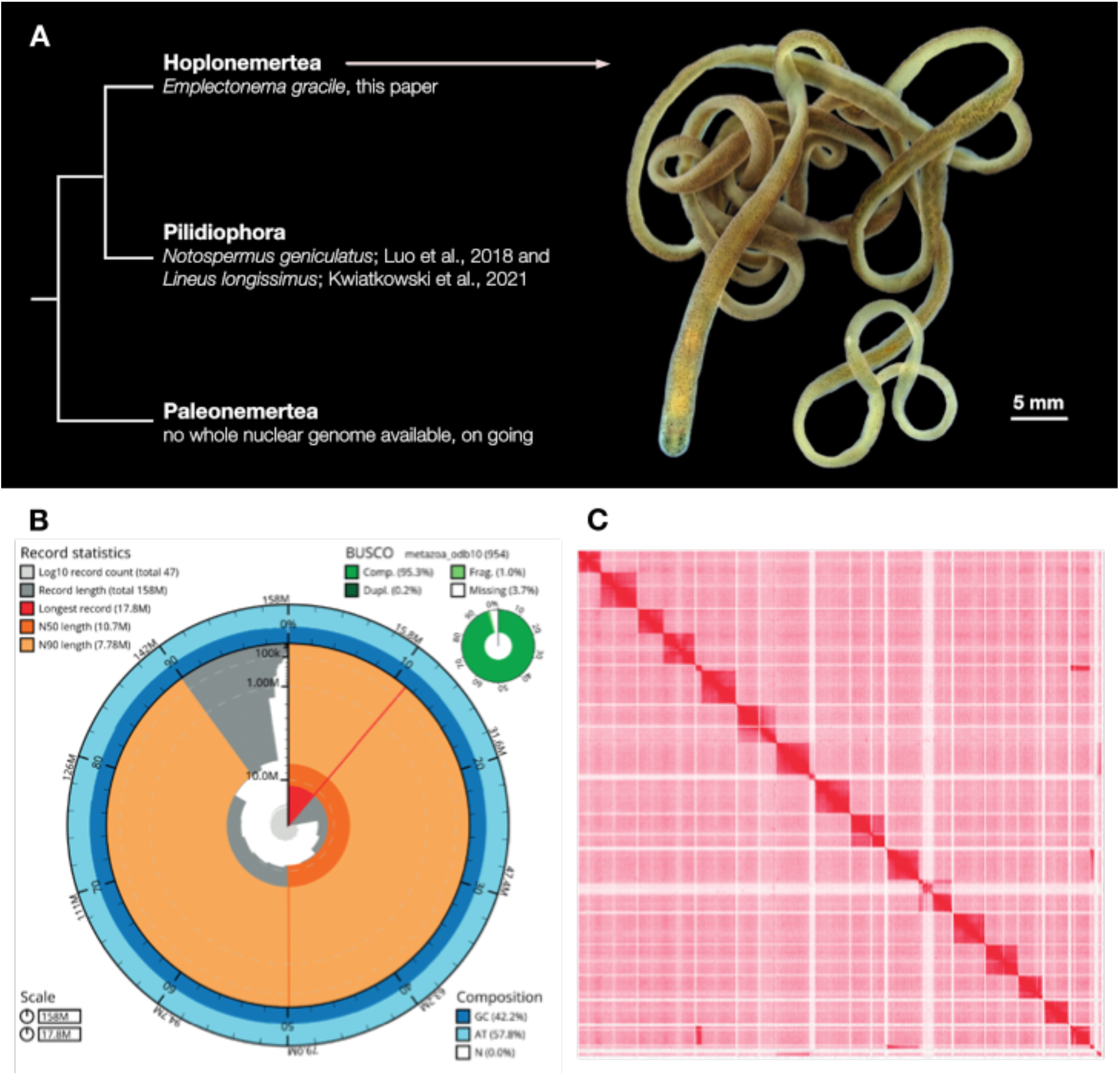
The genome of *Emplectonema gracile*. **(A)** Phylogeny of subgroups within the Nemertea phylum, with information about the species for which a whole genome is available (left), and adult specimen of *Emplectonema gracile* (right), photo taken by AVG. (**B)** “Snailplot” produced with BlobToolKit, illustrating N50 metrics and BUSCO gene completeness. **(C)** Hi-C contact map representing the final genome assembly of *E. gracile*, visualized with Juicebox 1.9.1.

In this study, we aimed to enrich the available genomic resources for Nemertea, a venomous animal group characterized by their regeneration capacities and obscure phylogenetic position (Stricker and Cloney 1983; Zattara et al. 2019). For it, we have sequenced, assembled, and annotated the genome of one representative of the Hoplonemertea clade, the species *Emplectonema gracile* (Figure 1A).

*Emplectonema gracile* inhabits the rocky shores of the North Atlantic Ocean and the Mediterranean Sea. This species has been selected for ease of collection, culturing, and spawning in the lab. Its slender, bi-toned body, armored with a venomous stylet used for capturing prey, can reach lengths of up to about 50 cm. The availability of the *E. gracile* genome will facilitate evolutionary, developmental, morphological, and toxicological studies within Nemertea, and will ultimately contribute to clarify the phylogenetic position of nemerteans within the Tree of life.

## Results and Discussion

For this diploid genome, HiFi sequencing yielded 26.6 Gbp of information contained in a total of 1,779,646 reads. Analysis of the genomic data with GenomeScope (Vurture et al. 2017) inferred a genome size of 157.9 Mbp with a heterozygosity of 1.5%, a uniqueness of 76.8%, and an error rate of 0.1% (Figure 1B, Figure S1A). K-mer analysis indicates the k-mers with the highest percentage out of all heterozygous k-mer pairs are diploid (92%) with only a small percentage of heterozygous k-mers being triploid or tetraploid (4%); these values indicate that the genome is diploid (Figure S1B).

Assembly, purging redundant haplotigs, and filtering out contamination resulted in a final genome assembly of 157.9 Mbp consisting of 22 contigs with an N50 of 10.0 Mbp (Figure 1B, Table 1). After scaffolding with Hi-C reads, the longest scaffold was 17.8 Mbp and the scaffold L50 was 6. The overall quality of the *E. gracile* genome was established by means of Mercury as 66.0 (Rhie et al. 2020). The assembly is rather complete with 95.3% of BUSCO markers detected (95.3% complete including 0.2% duplicated markers plus 1.0% fragmented). 99.8% of the k-mers mapped to the combined primary and alternative assembly, while 79.1% mapped to the assembly using only the primary one. The selected values for the different assembly parameters for each step (i.e., the unpurged primary genome assembly, the primary assembly after purging haplotypic duplications, the decontaminated primary genome assembly, and the HiC scaffolded genome) are shown in Table 1.

**Table 1.**
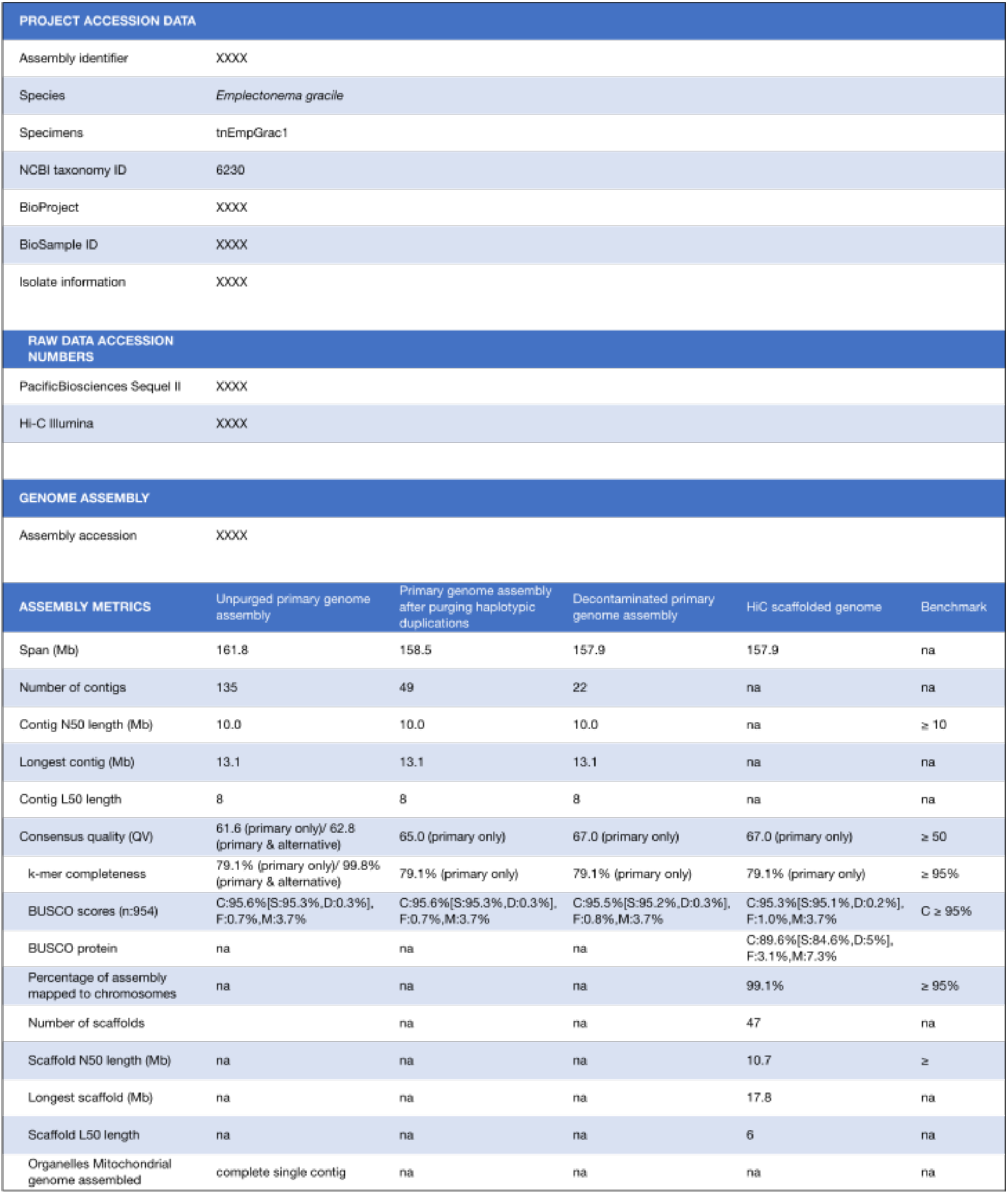
Project accession data and Assembly information for *E. gracile*. ^*^BUSCO scored based on the metazoan_odb10 BUSCO set using v5.1.2. C=complete [S=single copy, D=duplicated], F=fragmented, M=missing, n=number of orthologues in comparison.

| PROJECT ACCESSION DATA |  |  |  |  |  |
| --- | --- | --- | --- | --- | --- |
| Assembly identifier | XXXX |  |  |  |  |
| Species | Emplectonema gracile |  |  |  |  |
| Specimens | tnEmpGrac1 |  |  |  |  |
| NCBI taxonomy ID | 6230 |  |  |  |  |
| BioProject | XXXX |  |  |  |  |
| BioSample ID | XXXX |  |  |  |  |
| Isolate information | XXXX |  |  |  |  |
| RAW DATA ACCESSION NUMBERS |  |  |  |  |  |
| PacificBiosciences Sequel II | XXXX |  |  |  |  |
| Hi-C Illumina | XXXX |  |  |  |  |
| GENOME ASSEMBLY |  |  |  |  |  |
| Assembly accession | XXXX |  |  |  |  |
| ASSEMBLY METRICS | Unpurged primary genome assembly | Primary genome assembly after purging haplotypic duplications | Decontaminated primary genome assembly | HiC scaffolded genome | Benchmark |
| Span (Mb) | 161.8 | 158.5 | 157.9 | 157.9 | na |
| Number of contigs | 135 | 49 | 22 | na | na |
| Contig N50 length (Mb) | 10.0 | 10.0 | 10.0 | na | ≥ 10 |
| Longest contig (Mb) | 13.1 | 13.1 | 13.1 | na | na |
| Contig L50 length | 8 | 8 | 8 | na | na |
| Consensus quality (QV) | 61.6 (primary only)/ 62.8 (primary & alternative) | 65.0 (primary only) | 67.0 (primary only) | 67.0 (primary only) | ≥ 50 |
| k-mer completeness | 79.1% (primary only)/ 99.8% (primary & alternative) | 79.1% (primary only) | 79.1% (primary only) | 79.1% (primary only) | ≥ 95% |
| BUSCO scores (n:954) | C:95.6%[S:95.3%,D:0.3%], F:0.7%,M:3.7% | C:95.6%[S:95.3%,D:0.3%], F:0.7%,M:3.7% | C:95.5%[S:95.2%,D:0.3%], F:0.8%,M:3.7% | C:95.3%[S:95.1%,D:0.2%], F:1.0%,M:3.7% | C ≥ 95% |
| BUSCO protein | na | na | na | C:89.6%[S:84.6%,D:5%], F:3.1%,M:7.3% |  |
| Percentage of assembly mapped to chromosomes | na | na | na | 99.1% | ≥ 95% |
| Number of scaffolds |  | na | na | 47 | na |
| Scaffold N50 length (Mb) | na | na | na | 10.7 | ≥ |
| Longest scaffold (Mb) | na | na | na | 17.8 | na |
| Scaffold L50 length | na | na | na | 6 | na |
| Organelles Mitochondrial genome assembled | complete single contig | na | na | na | na |

For Hi-C, 52,305,505 reads were obtained. According to our Hi-C assembly and structural annotation, the *E. gracile* genome contains 15 putative chromosomes (Fig. 1C). After annotation, the resulting BUSCO protein score was 89.6% complete (84.6% single copy, 5% duplicated, 3.1% fragmented). This protein assessment was done employing the protein set BUSCO metazoan_odb10 dataset (Simão et al. 2015) combined with the selected nemertean transcriptomes shown in Table S1.

With values ranging between 157.9 and 161.8 Mbp, the *Emplectonema gracile* genome is shorter than the haploid size of 210 ± 5 Mbp previously estimated based on flow cytometry (Paule et al. 2021). Moreover, the *E. gracile* genome is substantially smaller than those of the two pilidiophoran species previously sequenced: *Lineus longissimus* (391 Mbp in 15 putative chromosomes distributed over 109 contigs) (Kwiatkowski et al. 2021), and *Notospermus geniculatus* (859 Mbp distributed over 60,228 contigs) (Luo et al. 2018).

## Materials and Methods

### Sampling

For this work, two specimens of *Emplectonema gracile* of unknown sex were selected. One was used for HiFi sequencing (specimen ID N59), while the other one was used for Hi-C sequencing (specimen ID N53) (Lieberman-Aiden et al. 2009; Hu et al. 2021). Both individuals were collected in Norway from the beach of Jeløya (Moss, Viken; N 59º25’23.6”, E 010º37’05.9”; WGS84; ±2 m). The specimens, approximately 10 cm long, were cut. The anterior body part of N53 was preserved in 4% formaldehyde as a voucher (Natural History Museum, University of Oslo, Norway; catalog number NHMO C7190). Remaining pieces of N53, and all N59, were flash frozen in liquid nitrogen.

### Identification

Morphological identification and DNA barcoding were performed for both specimens. Specimen N59 was barcoded using the DNA extracted for HiFi sequencing, while the DNA of N53 was extracted using the Qiagen DNeasy Blood and Tissue Kit following manufacturer’s instructions. 16S and 18S rRNA gene sequences of N59 were PCR amplified using the following primers: forward (16S: 5’ CCGGTCTGAACTCAGATCACGT 3’; 18S: 5’ CCCCGTAATTGGAATGAGTACA 3’), reverse (16S: 5’ CGCCTGTTTATCAAAAACAT 3’; 18S: 5’ AGCTCTCAATCTTGTCAATCCT 3’); and the following settings: 1x 2 minutes at 94 ºC, 40x [30 seconds at 94 ºC, 60 seconds at 51 ºC, 60 seconds at 72 ºC], 1x 2 minutes at 72 ºC. For N53, CO1 was PCR amplified using forward primer LCO1490-JJ (5’ CHACWAAYCATAAAGATARYGG 3’), and reverse primer HCO2198 JJ (5’ AWACTTCVGGRTGVCCAAARAARCA 3’) (Astrin and Stüben 2008). Resulting PCR products were Sanger sequenced by Macrogen (Amsterdam). The 16S (XXX), 18S (XXX), and COI (XXX) sequences confirmed the morphological identification. The COI sequence for N53 was identical to the COI sequence of the publicly available mitochondrial genome of *E. gracile* (NC_016952.1). The mitochondrial genome of the *Emplectonema gracile* specimen N59 determined by BLAST had a >99% similarity to the previously published mitochondrial genome for the same species (NCBI accession number JF727825).

### Genome Sequencing

For HiFi sequencing, DNA was extracted from posterior parts of N59. Samples were weighed and minced on dry ice followed by tissue disruption using a TissueRuptor II (QIAGEN, Germany) on its maximum settings for 10 seconds. High molecular weight (HMW) DNA was extracted using the Nanobind Tissue Big DNA kit (Circulomics Inc, USA). DNA quality and concentration were determined with a Nanodrop UV/Vis spectrophotometer (Thermo Fisher Scientific, USA), a Qubit BR dsDNA assay (Thermo Fisher Scientific), a pulsed field gel, and a Fragment Analyzer (Agilent, USA) run. Low molecular weight DNA was removed using the BluePippin (Sage Science, USA) system with High Pass Plus Gel cassettes. DNA was further purified and concentrated using the AMpure XP purification kit (Beckman Coulter, USA). A final concentration of 143 ng/µL in a volume of 75 µL was obtained. The library for HiFi circular consensus sequencing was constructed and sequenced on a SEQUEL II (Pacific Biosciences) platform at the Norwegian Sequencing Centre (Oslo, Norway).

For Hi-C sequencing, a library was prepared using the Arima High Coverage Hi-C+ Kit (Arima Genomics, USA). Specifically, the restriction enzymes used for Arima Hi-C 2.0 cut at the following recognition sites: ^GATC, G^ANTC, C^TNAG, T^TAA. For this reaction, ∼40 mg of disrupted tissue was used. Subsequently, a library was generated following the manufacturer’s instructions. For quality control, a combination of Qubit (Thermo Fisher Scientific, USA) and Fragment Analyzer (Agilent, USA) measurements, as well as a Kapa library quantification kit for Illumina libraries (Roche, Switzerland), were used. The final barcoded library was pooled on a quarter S4 flow cell in 2x150 bp paired-end mode on an Illumina NovaSeq sequencer (Illumina, USA). Hi-C library preparation and sequencing were done at the Norwegian Sequencing Centre (Oslo, Norway).

### Genome Profiling

Genome profiling steps were followed to assess k-mer frequencies within raw sequencing reads, and to estimate major genome characteristics such as genome size, heterozygosity, and repetitiveness. The k-mer distribution, with a k-mer size of 21, was generated using Jellyfish 2.3.0 using default settings. Based on this k-mer distribution, SmudgePlot 0.2.4 was run to test the ploidy of the genome (Marcais et al. 2012; Ranallo-Benavidez et al. 2020). GenomeScope 2.0 (Ranallo-Benavidez et al. 2020) with a k-mer size of 21, diploid level and a high-bound value of 1 million, was used to calculate the genome size, repetitiveness, and heterozygosity for a diploid genome using a combinatorial approach fitting a mathematical model to the k-mer distribution.

### De Novo Genome Assembly

Assembly of HiFi reads was carried out with Hifiasm 0.18.2 (Cheng et al. 2021), using default settings. Haplotypic duplications were purged with Purge_dups 1.4 (Guan et al. 2020). The primary assembly was checked for contamination and corrected using the BlobToolKit 3.1.4 software (Challis et al. 2020); all contigs not matching metazoan hits were excluded (Fig. S1C).

### Computational Scaffolding

The Arima Mapping Pipeline (https://github.com/ArimaGenomics/mapping_pipeline) was used for mapping raw Hi-C reads to the purged and decontaminated assembly outlined above. Briefly, Hi-C paired reads are first aligned to the reference independently using BWA-MEM to identify potential chimeric reads to be filtered out. Filtered single-end Hi-C reads are then paired and sorted based on mapping quality to produce a quality filtered, paired-end BAM file. Picard Tools is then used to flag PCR duplicates which are then discarded using SAMtools (Camacho et al. 2009). The quality filtered BAM file was then used as input for scaffolding. Scaffolding was performed in YaHS: yet another Hi-C scaffolding tool (https://github.com/c-zhou/yahs) (Zhou et al. 2022). The output of YaHS was then converted into .hic and .assembly files using Juicer tools 1.9.9_jcuda.0 (Durand et al. 2016) for manual curation in Juicebox Assembly Tools 1.9.1 (Dudchenko et al. 2018). The Hi-C contact map generation was done using Juicebox 1.9.1 (Robinson et al. 2018).

### Quality Control Checks

Several quality control checks were conducted after each analytical step (i.e., unpurged assembly, assembly after purging, decontaminated assembly, and scaffolded genome). Quast 5.0.2 (Gurevich et al. 2013) was used to determine statistical parameters of the primary genome assembly. Using Merqury 1.3 (Rhie et al. 2020), a meryl database was generated, and quality statistics such as consensus quality and k-mer completeness retrieved. The different assemblies were benchmarked against the 954 universal single-copy orthologs of the metazoa_odb10 dataset using BUSCO+ 5.5.0 (Simão et al. 2015, Manni et al. 2021). In preparation for BlobToolKit, Blast+ 2.13.0 (Camacho and Madden 2013) was used to map each contig of the assembly against a local copy of the NCBI nucleotide (nt) database downloaded as part of the pipeline. Additionally, HiFi reads were mapped against the primary assembly with Minimap2 2.17 (Li 2018), and further prepared for BlobTools with SAMtools 1.10 (Camacho et al. 2009).

The BUSCO scores, BLAST results, and read coverage were uploaded and further analyzed within BlobToolKit 3.1.4. After assembly, mitochondrial genome sequences were retrieved using Blast+ and the amino-acid sequences of all protein-coding genes of the mitochondrial genome NC_000931. These mitochondrial sequences were queried against the nt with Blast+ 2.13.0 to confirm species identification and possible sources of contamination.

### Genome Annotation

For structural annotation, RepeatModeler 2.0.1 (Flynn et al. 2020) was used to model repeat content followed by soft repeat masking utilizing RepeatMasker 4.1.2 with default settings (Smith et al. 2015). As no transcriptome was available for this species, annotation was performed using the Braker 3 (Hoff et al. 2019) pipeline based on protein sequences from closely related species. 37 publicly available nemertean transcriptomes belonging to 25 species (Table S1) were downloaded from NCBI, assembled with Trinity (Grabher et al. 2011), and translated with TransDecoder (https://github.com/TransDecoder/). The translated transcriptomes, combined with the OrthoDB v10 metazoa dataset, were used to generate protein prediction hints with ProtHint 2.6.0 (Hoff et al. 2019). The ProtHint mapping pipeline was used by Braker 3 to produce protein hints to train the model. The soft-masked and decontaminated primary genome assembly and protein databases were used as input to Braker 3. Gene annotations were assessed for completeness using BUSCO+ 5.5.0 (Simão et al. 2015) metazoa odb_10 database.

## Data Availability

European Nucleotide Archive: *Emplectonema gracile*. Accession number XXX. Unprocessed sequence data have been archived in the NCBI Sequence Read Archive under Bioproject XXX. The Refseq genome assembly can be found under accession XXX along with NCBI *E. gracile* Annotation release 1000. The genome sequence is released openly for reuse. All custom scripts are available at GitHub https://github.com/torstenstruck/InvertOmics and https://github.com/mkyapchiongco/Hi-C-WorkFlow.

## Ethical approval

The Nagoya protocol does not apply to this work. Both sample collection and molecular work were done in Norway.

## Acknowledgements

This work was funded by the Research Council of Norway project “InvertOmics – Phylogeny and evolution of lophotrochozoan invertebrates based on genomic data” (Project number: 300587 to THS). KMK, MKY, and NGR were funded by NSF DEB-1846174. We thank Matz Berggren (University of Gothenburg) for assisting and allowing AVG to use the camera setup used for taking the picture of a specimen of *E. gracile* included in Figure 1A, and Miguel Ángel Naranjo Ortiz (University of Oslo) and Emanuela Di Martino (University of Oslo) for discussion.

## Author Contributions

AVG and THS collected and preserved the specimens of *Emplectonema gracile*, identified by AVG. AVG carried out the molecular parts of this work which were not conducted at the Norwegian Sequencing Center, except for barcoding the voucher specimen N53, done by ATC. AVG ran the different genome profiling, genome assembly, and quality control checks developed by THS. MY-C performed Hi-C scaffolding of the genome plotted by AVG. NR performed the genome structural annotation. AVG, KMK, MM, and THS conceived the study. AVG wrote the first draft of the manuscript, and all authors approved the submitted version.

