## Supplementary Material for "First chromosome-level genome assembly of a ribbon worm from the Hoplonemertea clade, *Emplectonema gracile*, and its structural annotation"

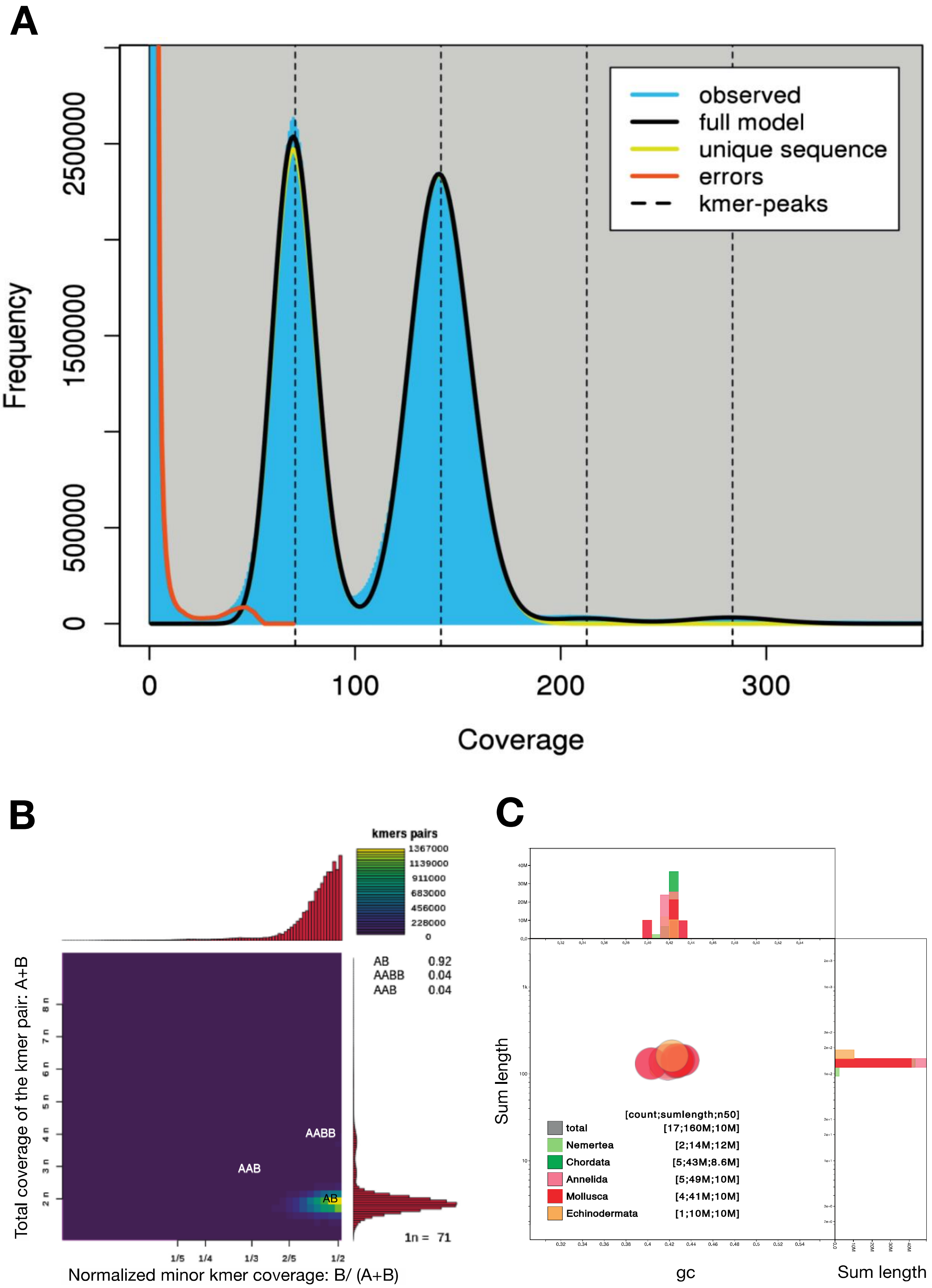

**Figure S1. –** (A) GenomeScope k-mer frequencies of the genome of *E. gracile* in its linear plot version. (B) Smudgeplot results presenting the estimated ploidy and genome structure of the *E. gracile* genome after analyzing the heterozygous k-mer pairs. (C) BlobToolKit plot showing the GC proportion and coverage levels of the genome produced.

| SPECIES | ACCESSION NUMB. | REFERENCE |
|---|---|---|
| *Amphiprous lactifloreus* | SRR11906528 | von Reumont et al. 2020 |
| *Antarctonemertes riesgoae* | SRR18913451 | Verdes et al. 2022 |
| *Antarctonemertes valida* | SRR18913444 | Verdes et al. 2022 |
| *Argonemertes australiensis* | SRR1506999 | Andrade et al. 2014 |
| *Argonemertes australiensis* | SRR1507001 | Andrade et al. 2014 |
| *Carinoma hamanako* | SRR1505092 | Andrade et al. 2014 |
| *Carinoma hamanako* | SRR1505094 | Andrade et al. 2014 |
| *Cephalothrix hongkongiensis* | SRR618505 | Andrade et al. 2014 |
| *Cephalothrix linearis* | SRR1275178 | Struck et al. 2014 |
| *Cephalothrix linearis* | SRR1273789 | Struck et al. 2014 |
| *Cephalothrix linearis* | SRR1273790 | Struck et al. 2014 |
| *Cephalothrix linearis* | SRR1275323 | Struck et al. 2014 |
| *Cephalothrix simula* | SRR18957873 | Vlasenko et al. 2022 |
| *Cephalothrix simula* | SRR18959724 | Vlasenko et al. 2022 |
| *Cerebratulus marginatus* | SRR618507 | Andrade et al. 2014 |
| *Cerebratulus* sp. | SRR1797867 | Egger et al. 2015 |
| *Lineus lacteus* | SRR1324979 | Ament-Velásquez et al. 2016 |
| *Lineus lacteus* | SRR1324980 | Ament-Velásquez et al. 2016 |
| *Lineus longissimus* | SRR1324981 | Ament-Velásquez et al. 2016 |
| *Lineus longissimus* | SRR1324986 | Ament-Velásquez et al. 2016 |
| *Lineus ruber* | SRR1324987 | Ament-Velásquez et al. 2016 |
| *Lineus ruber* | SRR1324988 | Ament-Velásquez et al. 2016 |
| *Lineus sanguineus* | SRR3581110 | Ament-Velásquez et al. 2016 |
| *Malacobdella grossa* | SRR1507002 | Andrade et al. 2014 |
| *Malacobdella grossa* | SRR1611560 | Kocot et al. 2017 |
| *Nipponnemertes* sp. | SRR1508368 | Andrade et al. 2014 |
| *Paranemertes peregrina* | SRR1507044 | Andrade et al. 2014 |
| *Paranemertes peregrina* | SRR1611562 | Kocot et al. 2017 |
| *Protopelagonemertes beebei* | SRR1507060 | Andrade et al. 2014 |
| *Riseriellus occultus* | SRR1505179 | Andrade et al. 2014 |
| *Riseriellus occultus* | SRR1505180 | Andrade et al. 2014 |
| *Tubulanus polymorphus* | SRR1611583 | Kocot et al. 2017 |
| *Tubulanus polymorphus* | SRR1273849 | Struck et al. 2014 |
| *Tubulanus polymorphus* | SRR1275406 | Struck et al. 2014 |
| *Tubulanus polymorphus* | SRR1273850 | Struck et al. 2014 |
| *Tubulanus polymorphus* | SRR1275407 | Struck et al. 2014 |
| *Tubulanus punctatus* | SRR1611583 | Andrade et al. 2014 |

**Table S1.** – List of transcriptomes used for annotation of the *Emplectonema gracile* genome.

| SOFTWARE TOOL | VERSION | SOURCE |
|---|---|---|
| **Augustus** | 3.3.3 | Hoff et al. 2019 |
| **Blast+** | 2.13.0 | Camacho et al. 2009 |
| **BlobToolKit** | 3.1.4 | Challis et al. 2020 |
| **Braker** | 3 | Stanke et al. 2006, Stanke et al. 2008, Hoff et al. 2016. Hoff et al. 2019; Bruna et al. 2021 |
| **BUSCO+** | 5.5.0 | Simão et al. 2015, Manni et al. 2021 |
| **GenomeScope** | 2.0 | Ranallo-Benavidez et al. 2020 |
| **HapPy** | 0.2.1 | Jiang et al. 2009 |
| **HifiAsm** | 0.18 | Cheng et al. 2021 |
| **Jellyfish** | 2.3.0 | Marcais et al. 2012 |
| **Juicebox assembly tools** | 1.9.1 | Robinson et al. 2018 |
| **Juicebox** | 1.11.08 | Dudchenko et al. 2018 |
| **Juicer tools** | 1.9.9 jcuda.0.8 | Durand et al. 2016 |
| **Merqury** | 1.3 | Rhie et al. 2020 |
| **Minimap2** | 2.17 | Li 2018 |
| **MitoHiFi** | 2.0 | Uliano-Silva et al. 2021 |
| **ProtHint** | 2.6.0 | Hoff et al. 2019 |
| **Purge_dups** | 1.4.2 | Guan et al. 2020 |
| **Quast** | 5.0.2 | Gurevich et al. 2013 |
| **RepeatMasker** | 4.1.6 | Smith et al. 2015 |
| **RepeatModeler** | 2.0.3 | Flynn et al. 2020 |
| **SAMtools** | 1.10 | Camacho et al. 2009 |
| **SmudgePlot** | 0.2.4 | Ranallo-Benavidez et al. 2020 |
| **Trinity** | 2.15.1 | Grabher et al. 2011 |
| **YAHS** | 1.2a.2 | Zhou et al. 2022 |

**Table S2.** – Software tools used.

**Supplementary Material References**


Ament-Velásquez SL, Figuet E, Ballenghien M, Zattara EE, Norenburg JL, Fernández-Álvarez FA, Bierne J, Bierne N, Galtier N. Population genomics of sexual and asexual lineages in fissiparous ribbon worms (*Lineus*, Nemertea): hybridization, polyploidy and the Meselson effect. Mol Ecol. 2016:25(14):3356–3369. https://doi.org/10.1111/mec.13717.

Andrade SC, Montenegro H, Strand M, Schwartz ML, Kajihara H, Norenburg JL, Turbeville JM, Sundberg P, Giribet G. A transcriptomic approach to ribbon worm systematics (Nemertea): resolving the Pilidiophora problem. Mol Biol Evol. 2014:31(12):3206–3215. https://doi.org/10.1093/molbev/msu253.

Brůna T, Hoff KJ, Lomsadze A, Stanke M, Borodovsky M. BRAKER2: automatic eukaryotic genome annotation with GeneMark-EP+ and AUGUSTUS supported by a protein database. NAR Genom Bioinform. 2021:3(1): lqaa108. https://doi.org/10.1093/nargab/lqaa108.

Camacho C, Coulouris G, Avagyan V, Ma N, Papadopoulos J, Bealer K, Madden TL. BLAST+: architecture and applications. BMC Bioinformatics. 2009:10:1–9. https://doi.org/10.1186/1471-2105-10-421.

Challis R, Richards E, Rajan J, Cochrane G, Blaxter M. BlobToolKit–interactive quality assessment of genome assemblies. G3 (Bethesda). 2020:10(4):1361–1374. https://doi.org/10.1534/g3.119.400908.

Cheng H, Concepcion GT, Feng X, Zhang H, Li H. Haplotype-resolved *de novo* assembly using phased assembly graphs with hifiasm. Nat Methods. 2021:18(2):170–175. https://doi.org/10.1038/s41592-020-01056-5.

Dudchenko O, Shamim MS, Batra SS, Durand NC, Musial NT, Mostofa R, Pham M, St Hilaire BG, Yao W, Stamenova E, Hoeger M, et al. The Juicebox Assembly Tools module facilitates de novo assembly of mammalian genomes with chromosome-length scaffolds for under $1000. BioRxiv. 2018: 254797. https://doi.org/10.1101/254797.

Durand NC, Shamim MS, Machol I, Rao SS, Huntley MH, Lander ES, Aiden EL. Juicer provides a one-click system for analyzing loop-resolution Hi-C experiments. Cell Syst. 2016:3(1):95–98. http://dx.doi.org/10.1016/j.cels.2016.07.002.

Egger B, Lapraz F, Tomiczek B, Müller S, Dessimoz C, Girstmair J, Škunca N, Rawlinson KA, Cameron CB, Beli E, et al. A transcriptomic-phylogenomic analysis of the evolutionary relationships of flatworms. Curr Biol. 2015:25(10):1347–1353. http://dx.doi.org/10.1016/j.cub.2015.03.034.

Flynn JM, Hubley R, Goubert C, Rosen J, Clark AG, Feschotte C, Smit AF. RepeatModeler2 for automated genomic discovery of transposable element families. Proc Natl Acad Sci USA. 2020:117(17): 9451–9457. https://doi.org/10.1073/pnas.1921046117.

Grabherr MG, Haas BJ, Yassour M, Levin JZ, Thompson DA, Amit I, Adiconis X, Fan L, Raychowdhury R, Zeng Q, et al. Trinity: reconstructing a full-length transcriptome without a genome from RNA-Seq data. Nat Biotechnol. 2011:29(7):644. http://dx.doi.org/10.1038/nbt.1883

Guan D, McCarthy SA, Wood J, Howe K, Wang Y, Durbin R. Identifying and removing haplotypic duplication in primary genome assemblies. Bioinformatics 2020:36(9):2896–2898. https://doi.org/10.1093/bioinformatics/btaa025.

Gurevich A, Saveliev V, Vyahhi N, Tesler G. QUAST: quality assessment tool for genome assemblies. Bioinformatics. 2013:29(8):1072–1075. https://doi.org/10.1093/bioinformatics/btt086.

Hoff KJ, Lange S, Lomsadze A, Borodovsky M, Stanke M. BRAKER1: unsupervised RNA-Seq-based genome annotation with GeneMark-ET and AUGUSTUS. Bioinformatics. 2016:32(5):767–769. https://doi.org/10.1093/bioinformatics/btv661.

Hoff KJ, Lomsadze A, Borodovsky M, Stanke M. Whole-genome annotation with BRAKER. *Gene Prediction: Methods and Protocols*. Humana, New York, NY. 2019, p. 65–95. https://doi.org/10.1007/978-1-4939-9173-0_5.

Hoff KJ, Stanke M. Predicting genes in single genomes with AUGUSTUS. Curr Protoc Bioinformatics. 2019:65(1):e57. https://doi.org/10.1002/cpbi.57.

Jiang Z, Rokhsar DS, Harland RM. Old can be new again: HAPPY whole genome sequencing, mapping and assembly. Int J Biol Sci. 2009:5(4):298. https://doi.org/10.7150/ijbs.5.298.

Kocot KM, Struck TH, Merkel J, Waits DS, Todt C, Brannock PM, Weese DA, Cannon JT, Moroz LL, Lieb B, et al. Phylogenomics of Lophotrochozoa with consideration of systematic error. Syst Biol. 2017:66(2):256–282. https://doi.org/10.1093/sysbio/syw079.

Li H. Minimap2: pairwise alignment for nucleotide sequences. *Bioinformatics.* 2018:34(18):3094–3100. https://doi.org/10.1093/bioinformatics/bty191.

Manni M, Berkeley MR, Seppey M, Simão FA, Zdobnov EM. BUSCO update: novel and streamlined workflows along with broader and deeper phylogenetic coverage for scoring of eukaryotic, prokaryotic, and viral genomes. MBE. 2021:38(10):4647–4654. https://doi.org/10.1093/molbev/msab199.

Marcais G, Kingsford C. Jellyfish: A fast k-mer counter. Version 1.1.4. Tutorialis e Manuais. 2012:1:1–8.

Ranallo-Benavidez TR, Jaron KS, Schatz MC. GenomeScope 2.0 and Smudgeplot for reference-free profiling of polyploid genomes. Nat Commun. 2020:11(1):1432. https://doi.org/10.1038/s41467-020-14998-3.

Rhie A, Walenz BP, Koren S, Phillippy AM. Merqury: reference-free quality, completeness, and phasing assessment for genome assemblies. Genome Biol. 2020:21(1):1–27. https://doi.org/10.1186/s13059-020-02134-9.

Robinson JT, Turner D, Durand NC, Thorvaldsdóttir H, Mesirov JP, Aiden EL. Juicebox. js provides a cloud-based visualization system for Hi-C data. Cell Systems. 2018:6(2): 256–258.

Simão FA, Waterhouse RM, Ioannidis P, Kriventseva EV, Zdobnov EM. BUSCO: assessing genome assembly and annotation completeness with single-copy orthologs. Bioinformatics. 2015:31(19):3210–3212. https://doi.org/10.1093/bioinformatics/btv351.

Smit AFA, Hubley R, Green P. RepeatMasker Open-4.0. 2015:2013–2015.

Stanke M, Diekhans M, Baertsch R, Haussler D. Using native and syntenically mapped cDNA alignments to improve de novo gene finding. Bioinformatics. 2008:24(5):637–644. https://doi.org/10.1093/bioinformatics/btn013.

Stanke M, Schöffmann O, Morgenstern B, Waack S. Gene prediction in eukaryotes with a generalized hidden Markov model that uses hints from external sources. BMC Bioinformatics. 2006:7(1):1–11. https://doi.org/10.1186/1471-2105-7-62.

Struck TH, Wey-Fabrizius AR, Golombek A, Hering L, Weigert A, Bleidorn C, Klebow S, Iakovenko N, Hausdorf B, Petersen M, et al. Platyzoan paraphyly based on phylogenomic data supports a noncoelomate ancestry of Spiralia. MBE. 2014:31(7):1833–1849. https://doi.org/10.1093/molbev/msu143.

Uliano-Silva M, Ferreira JGR, Krasheninnikova K, Formenti G, Abueg L, Torrance J, Myers EW, Durbin R, Blaxter M, McCarthy SA. MitoHiFi: a python pipeline for mitochondrial genome assembly from PacBio high fidelity reads. BMC Bioinformatics. 2023:24(1):288. https://doi.org/10.1186/s12859-023-05385-y.

Verdes A, Taboada S, Hamilton BR, Undheim EAB, Sonoda GG, Andrade SCS, Morato E, Marina AI, Cárdenas CA, Riesgo A. Evolution, expression patterns, and distribution of novel ribbon worm predatory and defensive toxins. Mol Biol Evol. 2022:39(5):msac096. doi: 10.1093/molbev/msac096.

Vlasenko AE, Kuznetsov VG, Magarlamov TY. Investigation of peptide toxin diversity in ribbon worms (Nemertea) using a transcriptomic approach. Toxins. 2022:14(8):542. https://doi.org/10.3390/toxins14080542.

von Reumont BM, Lüddecke T, Timm T, Lochnit G, Vilcinskas A, von Döhren J, Nilsson MA. Proteo-transcriptomic analysis identifies potential novel toxins secreted by the predatory, prey-piercing ribbon worm *Amphiporus lactifloreus*. Mar Drugs. 2020:18(8):407. https://doi.org/10.3390/md18080407.

Zhou C, McCarthy SA, Durbin R. YaHS: yet another Hi-C scaffolding tool. Bioinformatics. 2023:39(1):btac808. https://doi.org/10.1093/bioinformatics/btac808.
